# Development and Characterization of Ultrasound-Activated Polymeric Microdroplets for Targeted Chemotherapy

**DOI:** 10.64898/2026.06.28.735147

**Authors:** Joshua Antonio Whiting, Audri Yasmin Al Hasan Dara, James Francis Kwan, Jan Kubanek

## Abstract

Potent antineoplastics, such as afatinib and freebase doxorubicin, are associated with systemic toxicity. To address this issue, we developed a carrier that releases drugs, including afatinib and doxorubicin, specifically at the focus of low-intensity ultrasound. This remotely triggered and focal approach enables the release of drugs specifically at the ultrasound focus, thus mitigating undesirable off-target effects, and at concentrations governed by the duration of the applied ultrasound. We produced ultrasound-sensitive microdroplets with high encapsulation efficiencies (39.6% for afatinib and 46.6% for doxorubicin). The microdroplets consist of an ultrasound-sensitive drug delivery system based on a methoxy poly(ethylene glycol)-poly(D, L-lactide) diblock copolymer (mPEG-PDLLA) and perfluorooctyl bromide (PFOB). Antineoplastic agents were encapsulated within these microdroplets via co-evaporation during particle synthesis. The microdroplets released doxorubicin and afatinib in an ultrasound-pressure-dependent manner, with fitted half-maximal release pressures (*P*_50_) of 0.61 MPa and 0.72 MPa, respectively. Together, the effective encapsulation of hydrophobic antineoplastic agents and the dose-dependent ultrasound-triggered release provide a new method for targeted drug delivery and a foundation for future targeted chemotherapies.

**Graphical Abstract:** 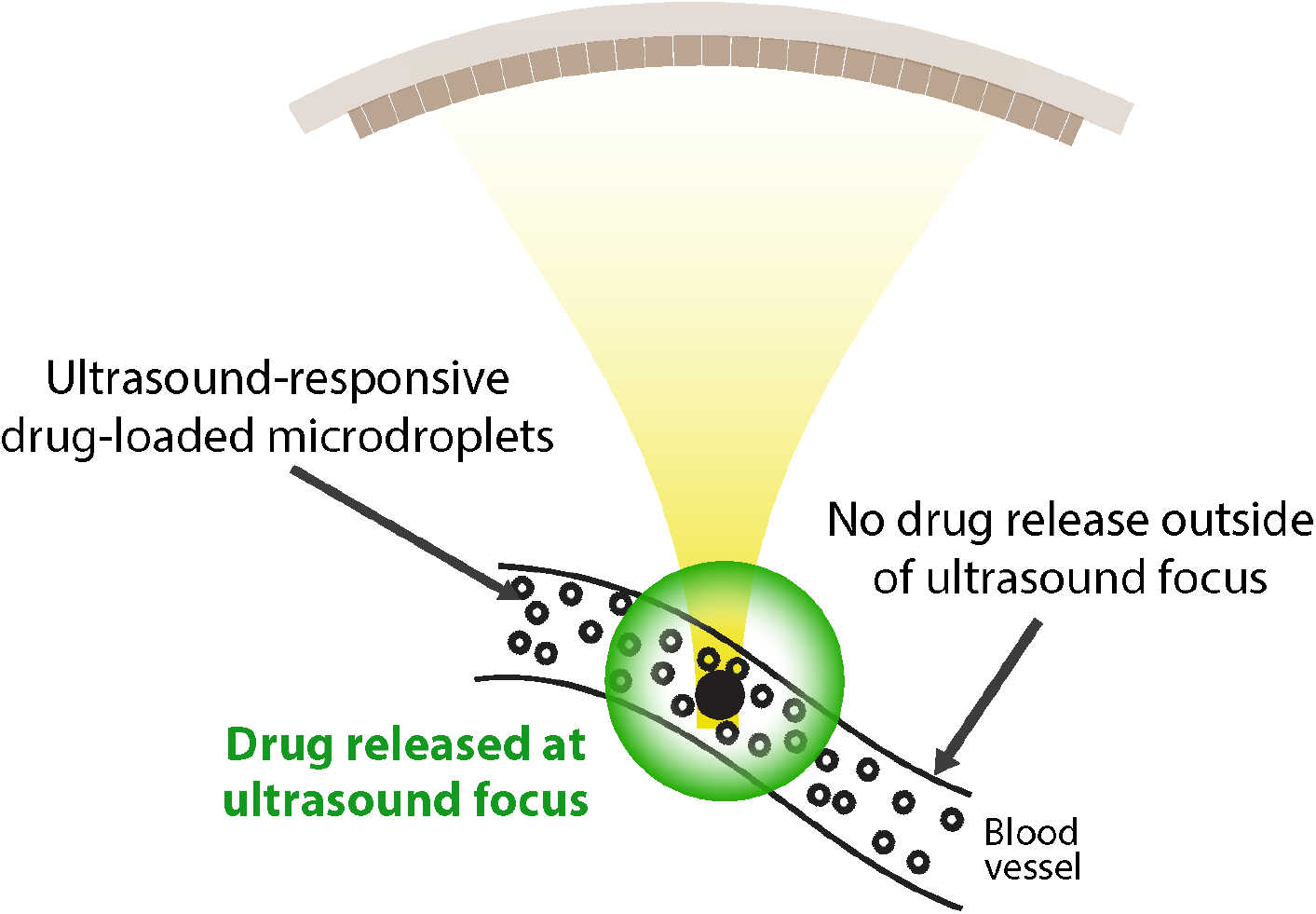

**Highlights:**

- We developed an ultrasound-triggered drug release vehicle using biocompatible materials.
- The vehicle has high encapsulation efficiency for hydrophobic chemotherapeutics.
- We demonstrated dose-dependent release.
- The vehicle provides a platform for targeted chemotherapy.

## 1. Introduction

Glioblastoma Multiforme (GBM) is the most aggressive and lethal primary brain tumor. Existing therapies, which include surgical resection, temozolomide chemotherapy, and radiotherapy can provide only limited increase in life expectancy, which averages to about 15 months following diagnosis. This poor prognosis is due to the cancer cells’ rapid proliferation, diffuse infiltration, and high degree of therapeutic resistance [1]. A significant barrier to effective treatment is the blood-brain barrier (BBB), a highly selective endothelial barrier that severely restricts the passage of most systemically administered antineoplastic agents from the bloodstream into the brain parenchyma [2]. Potent antineoplastic agents such as freebase doxorubicin (DOX), a hydrophobic form of the drug, and afatinib exhibit strong cytotoxic activity and inhibition against a broad range of cancers, including GBM. Still, their clinical utility is severely limited by the systemic toxicity and associated adverse side effects [3, 4]. Moreover, the drugs suffer from poor solubility, rapid systemic clearance, and inadequate penetration across the BBB [5, 6]. Consequently, there is a pressing need for a delivery platform that can protect these drugs from interacting with the body systemically, bypass the BBB, and release them on demand directly within the tumor microenvironment. Overcoming these challenges will expand the utility of this delivery system to many cancers and conditions that can benefit from targeted DNA damage and manipulation of proliferation signaling.

Focused ultrasound (fUS) enables local drug delivery at the ultrasound focus, sparing other tissues and organs from potential harmful side effects [7, 8]. Signif-icant research has been dedicated to the development of ultrasound-responsive carriers for controlled drug release. For example, DOX and its salt form, Doxorubicin-HCl, have been encapsulated in various systems, including PLGA nanoparticles [9, 10], liposomes [11], and hydrogel nanoparticles [12]. These systems have demonstrated the feasibility of ultrasound-triggered release and improved efficacy over systemic delivery in preclinical models [13, 14]. Despite these advances, many existing platforms are optimized for hydrophilic or moderately soluble drugs and exhibit limited encapsulation efficiency for highly hydrophobic antineoplastics [15]. In addition, several systems rely on post-loading strategies or exhibit inconsistent pressure-dependent release, limiting their ability to achieve predictable, on-demand drug delivery.

Our group has previously deployed in non-human primates (NHPs) an ultrasound sensitive drug delivery platform based on a methoxy poly(ethylene glycol)-poly(D,L-lactide) (mPEG-PDLLA) diblock copolymer. These nanoparticles can safely and selectively release propofol in deep brain regions of NHPs upon the trigger of transcranial focused ultrasound [16, 17]. The nanoparticles showed an excellent safety profile, with repeated applications well tolerated by NHPs [16, 17].

Building upon this validated platform, in this work, we developed a co-evaporation-based synthesis method to efficiently encapsulate low-solubility hydrophobic agents. This co-evaporation approach promotes mixing between the hydrophobic PDLLA block of the polymer and the drug molecules during solvent removal, thereby enhancing encapsulation of poorly soluble compounds. This contrasts traditional post-loading methods [16, 17], which cannot effectively encapsulate these insoluble hydrophobic compounds. Together, this work provides a prescription for the development of ultrasound-sensitive drug carriers that can effectively encapsulate and release hydrophobic compounds, with special focus on antineoplastic drugs.

## 2. Materials and Methods

### 2.1. Materials

MPEG-PDLLA diblock copolymer with a molecular weight ratio of 2,000:2,200 g/mol was procured from PolySciTech. The active antineoplastic agents, afatinib and DOX, were sourced from AdooQ Bioscience and MedKoo Bioscience, respectively. Additionally, the ultrasound-sensitive agent, PFOB, was sourced from Sigma-Aldrich. All solvents, including HPLC-grade tetrahydrofuran (THF) and methanol, were obtained from Fisher Scientific. Phosphate-buffered saline (PBS, 1X) was purchased from Gibco. Emulsification was performed using a 20 kHz, 500 W probe sonicator (VCX500, Sonics & Materials Inc.). Ultrasound stimuli were generated by a function generator (33520b, Keysight) and amplified through a 55-dB power amplifier (A150, Electronics & Innovation) with an operational range of 300 kHz–30 MHz.

### 2.2. Synthesis of Drug-Loaded Copolymer Microdroplets

The synthesis of drug-loaded copolymer microdroplets was performed using a modified co-evaporation protocol (see Fig. 1) designed to maximize the hydrophobic interaction between the poly (lactic acid) block of the copolymer and our antineoplastic agents, ensuring high encapsulation efficiency. Briefly, 16 mg of MPEG-PDLLA copolymer was dissolved in 1 mL of THF within a 1.5 mL microcentrifuge tube. 80 µL of a 0.1 g/mL solution of either afatinib or DOX in Dimethyl Sulfoxide (DMSO) was added to the polymer/THF mixture and vortexed to ensure complete homogenization. The tube was left uncapped overnight in a room-temperature fume hood to allow for complete evaporation of the organic solvent, resulting in a polymer-drug film deposited on the walls and bottom of the tube. Residual solvent was removed under vacuum as needed prior to rehydration. Subsequently, 1 mL of PBS was added to the polymer-drug film and left on a shaker for 30 minutes to facilitate rehydration and dispersion. Subsequently, 48 µL of PFOB and the rehydrated solution were added to a 15 mL conical tube before the total volume was adjusted to 8 mL with PBS. The final mixture was emulsified using a 500 W probe sonicator operated at 20% amplitude with constant water circulation from an ice bath to maintain a consistent 10 °C. Three rounds of sonication were applied in 90-second intervals for a total duration of 4.5 minutes, yielding a stable emulsion of PFOB-integrated, drug-loaded copolymer microdroplets. These final microdroplets were then washed five times by centrifugation (5000 rpm, 5 minutes, 4 °C) to pellet the intact microdroplets; the supernatant, containing free polymer and unencapsulated drug, was discarded before the pellet was resuspended in fresh PBS for further washing or analysis.

**Figure 1:**
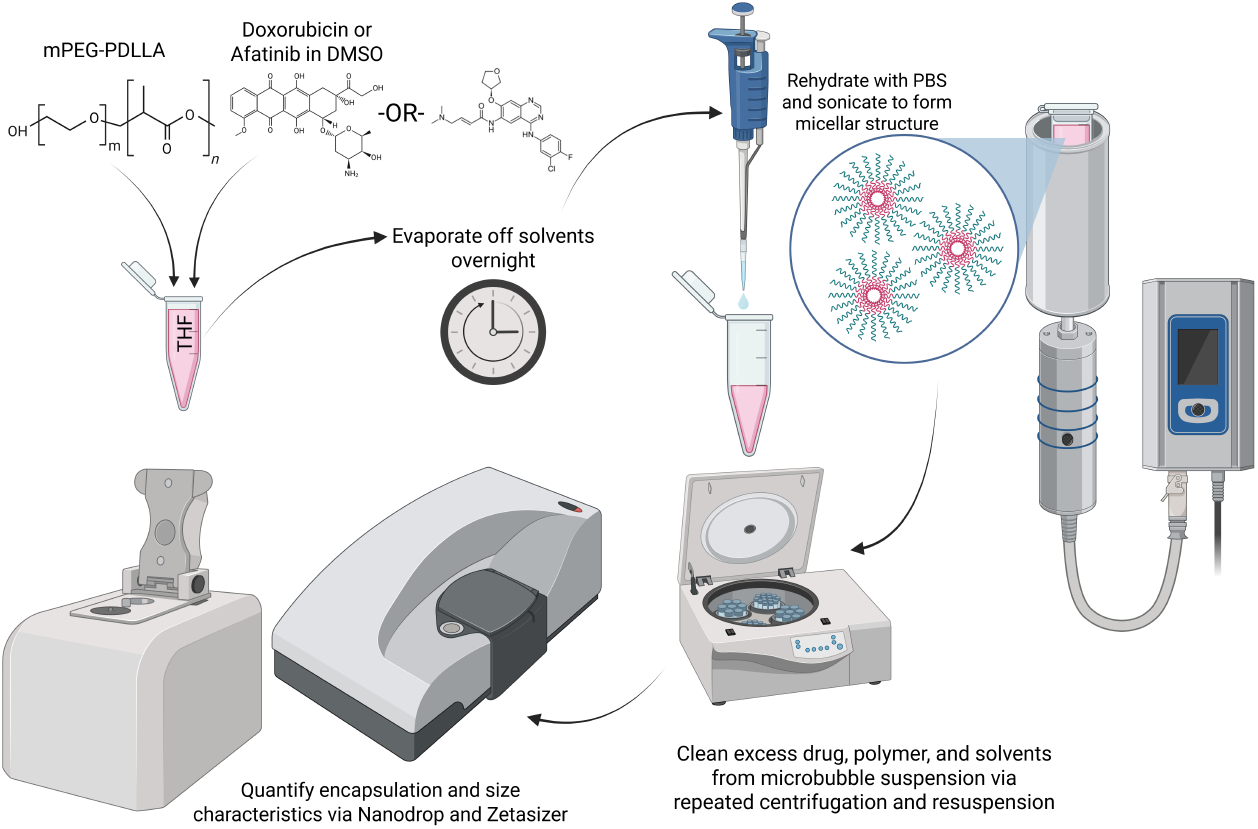
Schematic illustration of the co-evaporation synthesis process for drug-loaded mPEG–PDLLA microdroplets. (A) Dissolution of copolymer and drug in THF, (B) solvent evaporation to form a polymer–drug film, (C) rehydration with PBS then addition of PFOB, and (D) bath sonication to generate stable ultrasound-sensitive microdroplets.

### 2.3. Characterization of Copolymer Microdroplets

The hydrodynamic diameter and size distribution of the prepared micro-droplets were determined via dynamic light scattering (DLS) using a Zetasizer Nano S (Malvern Panalytical, UK). Measurements were performed on undiluted suspensions at 25 °C, and reported values represent the mean ± standard deviation of the intensity-weighted distribution across three measurements, each consisting of 17 runs. Due to the multimodal size distribution of the micro-droplets, polydispersity index (PDI) values were not reported, as PDI assumes a unimodal distribution. Instead, size distributions were interpreted qualitatively with scanning electron microscopy (SEM). SEM was done using a FEI Quanta 600 F scanning electron microscope at 5 kV to 10 kV. Microdroplet samples were deposited onto silicon wafer–topped SEM stubs, dried under nitrogen, sputter-coated with a thin (10-20 nm) layer of gold-palladium to reduce charging artifacts, and imaged by SEM.

Encapsulation efficiency (EE) was quantified by vortexing a 10 µL aliquot of the microdroplet solution with 90 µL of methanol to disrupt the copolymer shell and liberate the encapsulated drug. The concentration of the drug in methanol was then determined using a UV-Vis spectrophotometer (NanoDrop 2000, Thermo Scientific). Absorbance was measured in triplicate for each sample at a wavelength of 343 nm for afatinib and 495 nm for DOX, with concentrations calculated against standard curves of the pure drugs in methanol. Linearity was ensured throughout all relevant concentration ranges measured. The EE was calculated as follows:

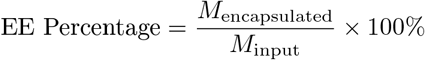

Additionally, the mass of the drug in the waste was measured to account for the total starting mass of the drug added to the synthesis system, and was confirmed to be within a 90-100% recovery range. Concentrations in mg/ml were found based on standard curves with the following equations:

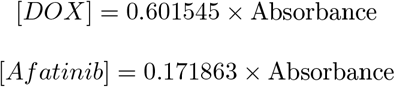

Concentrations were converted to masses by multiplying by the volume of the microdroplet solution and waste, respectively. Subsequent recovery percentages were then found for each antineoplastic agent with the following equation:

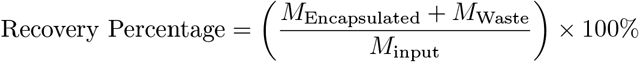

**Table 1:** Average microdroplet parameters. Size and Encapsulation are presented as Average ± Standard Deviation evaluated over the fifty most recent microdroplet syntheses (n=50). Release presented as Average ± Standard Deviation evaluated over three independently synthesized batches, with each measurement conducted in triplicate (n=9).

| Agent | Average Size<br>($\mu\text{m}$) | Average<br>EE (%) | Release<br>Percentage<br>(1.3 MPa) | Mass<br>Recovery<br>(%) |
| --- | --- | --- | --- | --- |
| **Doxorubicin** | 2.22 ± 0.77 | 46.6 ± 9.8 | 47.28 ± 3.19 | 90 |
| **Afatinib** | 1.31 ± 0.48 | 39.6 ± 10.7 | 45.35 ± 1.28 | 98 |

### 2.4. In Vitro Ultrasound-Triggered Drug Release

The ultrasound-triggered release profile was characterized using an in vitro setup. Briefly, 250 µL of freshly synthesized DOX-loaded microdroplets were placed in a standard 1.5 mL microcentrifuge tube. To this tube, 250 µL Ethyl Acetate (EA) was added to act as a hydrophobic sink to capture the released drug. EA was selected for the sink as it did not disrupt the microdroplet shell without sonication, was well separated from the microdroplet solution, and exhibited acceptable solubility of both DOX and afatinib. This tube was positioned in a holder in a degassed water bath, and a 300 kHz focused ultrasonic transducer (H-115, Sonic Concepts) was placed below the sample holder at the focal distance. The stimulation protocol consisted of continuous 10-ms pulses delivered at a rate of 10 pulses per second for a total sonication duration of 60 seconds. These stimulation parameters were used throughout all experimentation, with amplitude adjusted to modulate pressure. The pressure amplitude was systematically varied between 0 and 1.8 MPa (0, 0.3, 0.6, 0.9, 1.1, 1.3, 1.5, 1.8 MPa – see Fig. 2). These conditions were selected to probe the pressure-dependent activation of the microdroplets. Acoustic pressures reported correspond to peak negative pressures measured at the focal point in degassed water using a calibrated fiber-optic hydrophone (Precision Acoustics Ltd.). After sonication and a total incubation time of 2 minutes, 50 µL of the EA layer was carefully extracted, avoiding the microdroplet aqueous layers. The concentration of DOX partitioned into the ethyl acetate was quantified via UV-Vis spectrophotometry, and the percentage of drug released was calculated relative to the total encapsulated amount determined for that batch.

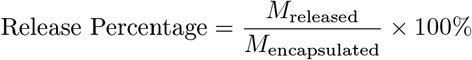

**Figure 2:**
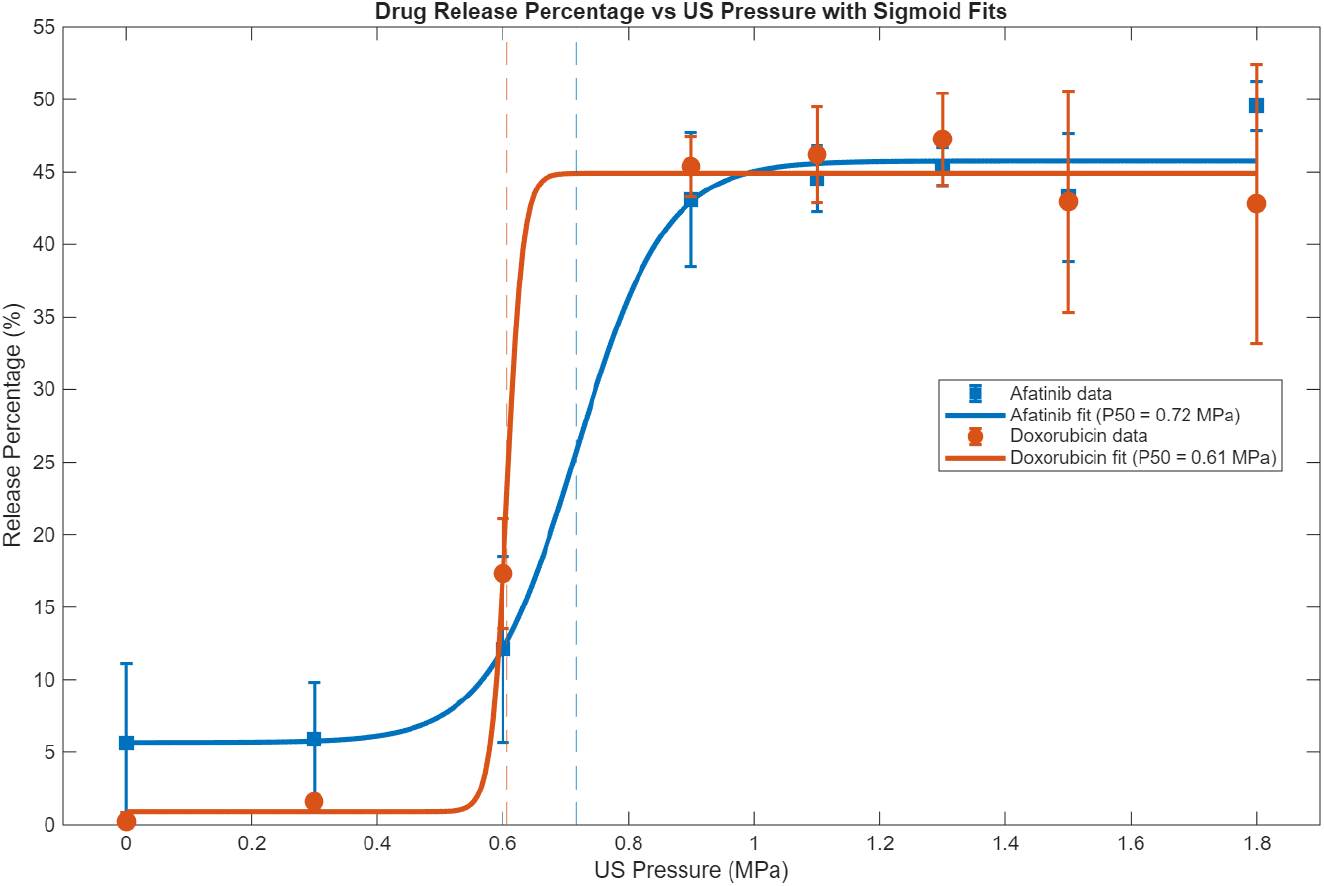
Ultrasound-triggered release of doxorubicin (red) and afatinib (blue) from mPEG–PDLLA microdroplets as a function of applied acoustic pressure amplitude. Data were fit using a four-parameter logistic model. The fitted half-maximal release pressures (*P*_50_) were 0.61 MPa for doxorubicin and 0.72 MPa for afatinib (*R*^2^ = 0.994 and 0.992, respectively). Data represent mean ± SD (n = 3) normalized to the encapsulated drug concentration for each batch measured in triplicate.

The same method was used to quantify afatinib release, with slight modifications – an increased total incubation time of 6 minutes was employed to allow for a 3-minute centrifugation step, which helped separate the organic and aqueous layers after sonication. Conversion from absorbance to concentration in mg/ml was done based on standard curves using the following equations:

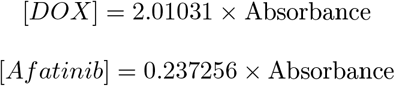

Analysis of the short-term release profile confirmed that nearly all release occurs during sonication, and that additional incubation does not result in greater release levels. Additionally, control experiments confirmed that the centrifugation step itself did not induce additional release, ensuring that the release profiles for both drugs are directly comparable.

Release-pressure data were fit using a four-parameter logistic model

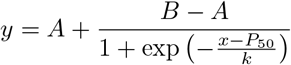

where *A* and *B* are the lower and upper asymptotes, *P*_50_ is the pressure producing half-maximal release, and *k* is the slope parameter.

## Results

### 3.1. Microdroplet Synthesis and Encapsulation Efficiency

For effective ultrasound-based release of hydrophobic drugs from the micro-droplets, we have developed a co-evaporation synthesis technique (see Fig. 1). The technique proved effective for generating copolymer microdroplets with high drug loading capabilities. Quantitative analysis using UV-Vis spectrophotometry consistently revealed high EEs across multiple batches. The EE for DOX was 46.56% ± 9.8% (mean ± SD), while afatinib had an encapsulation efficiency of 39.57% ± 10.67%. The relatively high encapsulation efficiencies are attributed to strong hydrophobic interactions between the drug molecules and the PDLLA block during co-evaporation, which promotes drug retention within the polymer domain of the microdroplets. These encapsulation efficiencies demonstrate effective loading of both antineoplastic agents and are comparable to or exceed values reported for similar polymer-based delivery systems [12, 10, 15]. These measurements were further supported by mass balance analysis, which consistently demonstrated >90% recovery across formulations.

### 3.2. Ultrasound-Triggered Drug Release

Our previous studies in NHPs [16, 17] demonstrated that the incorporation of PFOB confers ultrasound sensitivity to the microdroplets. In this study, we re-evaluated this capacity given the new co-evaporation method. Indeed, the release of encapsulated drugs was found to be critically dependent on the applied ultrasound pressure amplitude. Sigmoidal fitting of the pressure-response curves yielded half-maximal release pressures (*P*_50_) of 0.61 MPa for doxorubicin and 0.72 MPa for afatinib, indicating that substantial drug release can be achieved at relatively low acoustic pressures. Release increased rapidly between approximately 0.6 and 0.9 MPa and reached a plateau near 1.3 MPa, suggesting saturation of microdroplet activation at higher pressures. The observed threshold behavior is consistent with ultrasound-induced mechanical disruption of the microdroplet shell, which may include cavitation-related effects, acoustic radiation forces, and/or local shear. Notably, both threshold pressures are well below the FDA diagnostic ultrasound guideline corresponding to a mechanical index less than 1.9 [18]. Although the *in-vitro* release system used here does not fully recapitulate the hemodynamic conditions of our previous NHP studies, the findings are consistent with those previous studies [16, 17]. Therefore, this maximal plateau pressure level was adopted for all subsequent release characterization experiments.

### 3.3. Size and Stability

Dynamic light scattering and scanning electron microscopy analysis revealed that the co-evaporation method yielded microdroplets with a mean hydrodynamic diameter of approximately 2.2 µm for DOX particles, and a mean hydrodynamic diameter of approximately 1.3 µm for Afatinib particles. While the micron-scale size of these carriers could limit deep vascular penetration, it may be advantageous for intravascular ultrasound-mediated drug release, where transient trapping within tumor microvasculature can enhance local drug concentration. Moreover, a larger nanoparticle size can encapsulate substantially larger volume of encapsulated drug.

Scanning electron microscopy (SEM) provided complementary confirmation of microdroplet size and primarily spherical morphology, as shown in Fig. 3. Observed particle diameters were consistent with hydrodynamic sizes measured by dynamic light scattering, supporting the validity of the reported average diameters. Qualitative assessment of the SEM images indicated a moderately dispersed population, with the majority of microdroplets falling within approximately 1 µm of the mean diameter for each formulation. Occasional larger microdroplets were also observed, contributing to a broadened size distribution. This heterogeneity is consistent with the multimodal distributions observed in the dynamic light scattering measurements, and likely contributes to the inability to accurately report polydispersity index, as standard PDI analysis assumes a unimodal distribution.

**Figure 3:**
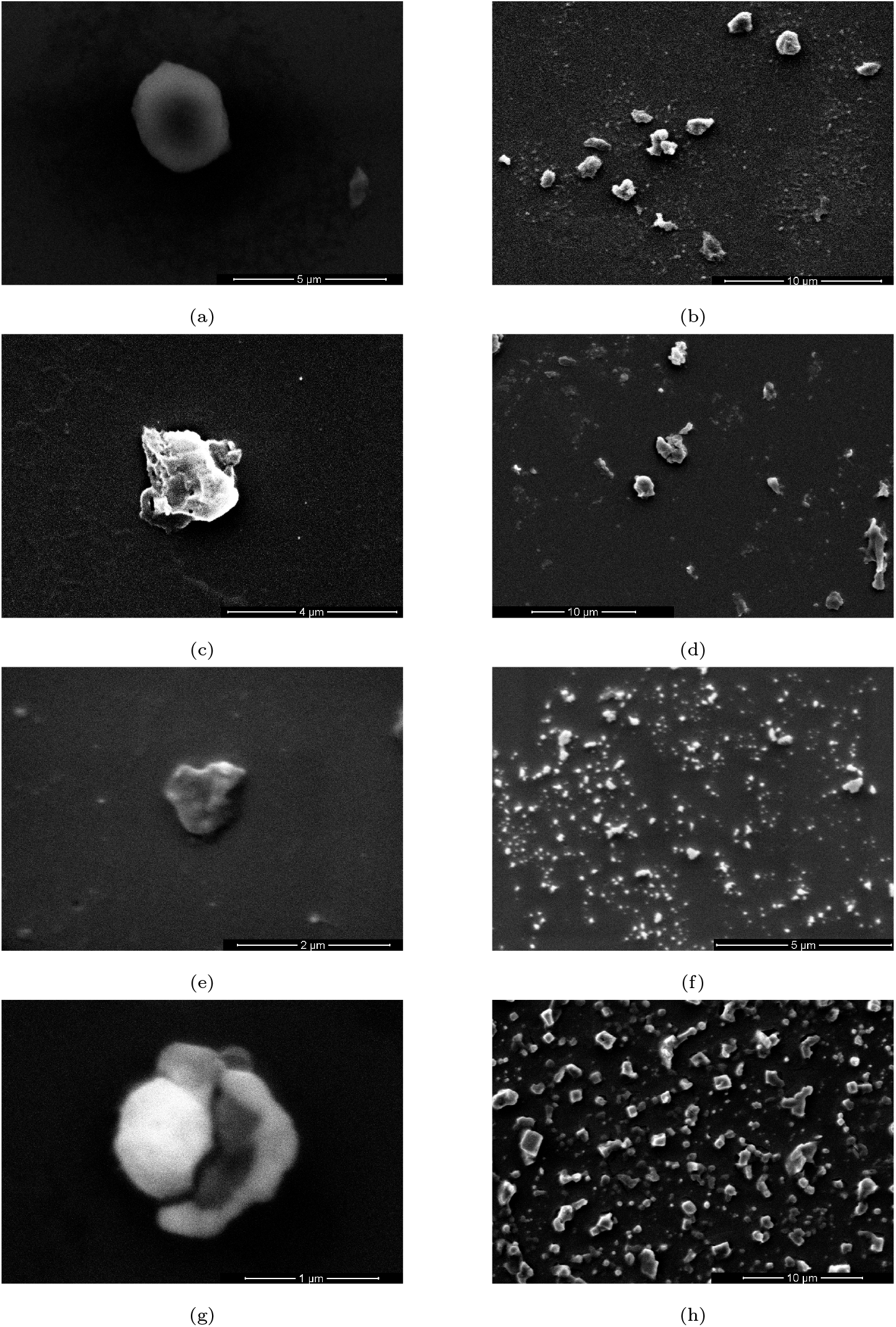
Scanning electron microscopy (SEM) images of mPEG–PDLLA microdroplets. (a,b) -loaded microdroplets showing a representative single particle and a population view. (c,d) Afatinib-loaded microdroplets. (e,f) Propofol-loaded nanodroplets. (g,h) Empty microdroplets.

The stability of the encapsulated carriers and their readiness to release their payload were assessed over 200 days, as shown in Fig. 4. During this assessment, microdroplets were stored in PBS in sealed centrifuge tubes at 4°C, protected from light. A gradual decrease in encapsulation efficiency was observed over time, consistent with drug leakage as the biodegradable copolymer matrix undergoes hydrolysis. This behavior is a critical parameter for defining the formulation’s usable shelf life. This leakage plateaued at approximately 20%, after which encapsulation efficiency remained stable for up to 38 weeks. This suggests that extended storage is possible as long as the microdroplets undergo additional cleaning and resuspension steps to eliminate the leaked drug. Release profiles did not appreciably change over time, as drug release scaled linearly with the remaining encapsulated drug at each time point.

**Figure 4:**
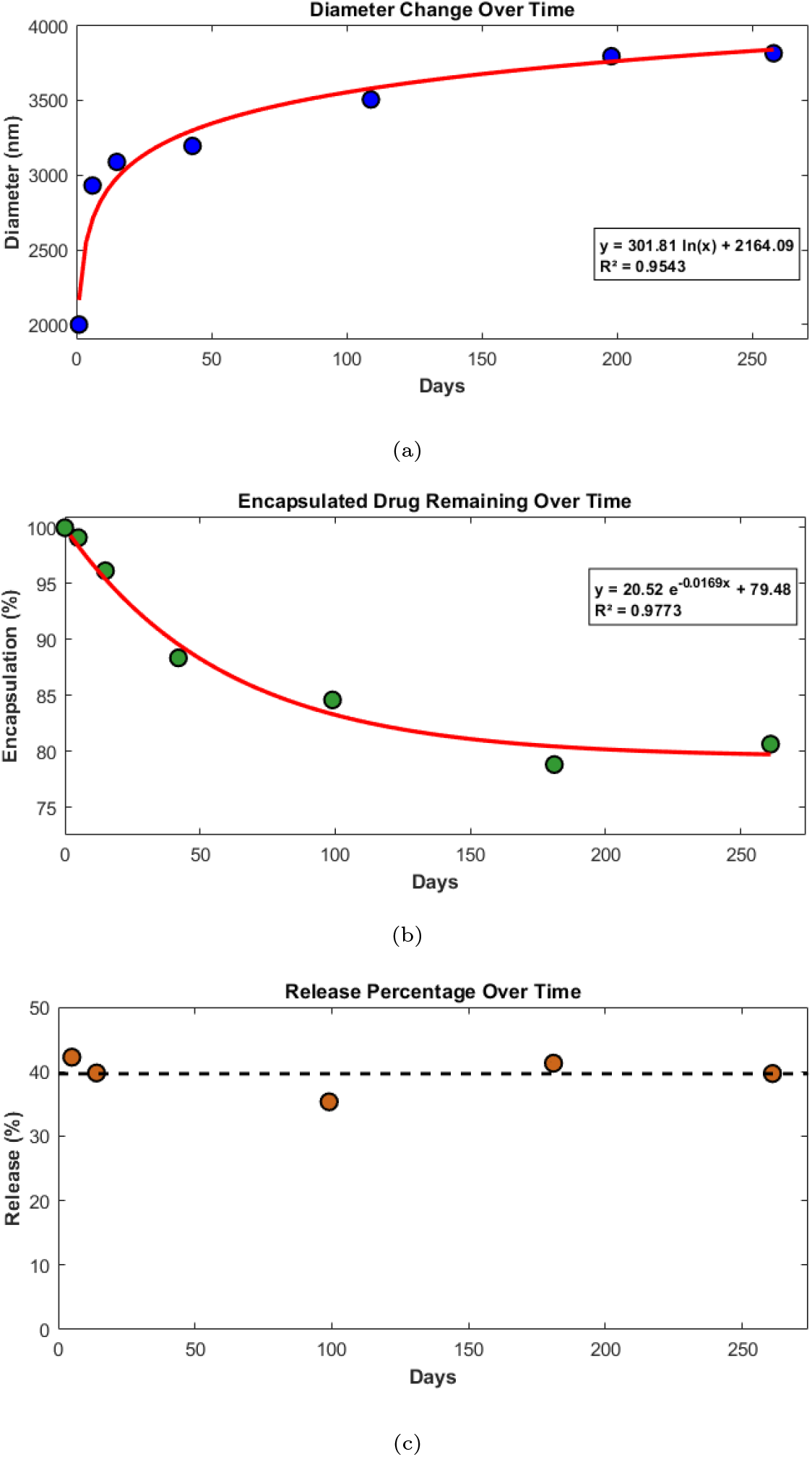
Long-term stability of drug-loaded microdroplets during storage. (a) Encapsulation efficiency over time, showing gradual drug leakage and stabilization at approximately 80% retention. (b) Hydrodynamic diameter measured by dynamic light scattering, indicating progressive aggregation during storage, reaching approximately 4 µm. (c) Ultrasound-triggered release at 1.3 MPa normalized to remaining encapsulated drug, demonstrating preserved release behavior over time.

Extended dynamic light scattering data also indicate an increase in the average hydrodynamic diameter, approaching 4 µm, throughout storage. The observed increase in hydrodynamic diameter over time is consistent with gradual microdroplet aggregation. Notably, aggregation appeared partially reversible via homogenization, suggesting that particle–particle interactions are not permanently fused but may be mitigated through mechanical disruption. Extended storage may therefore necessitate alternative methods of revival, such as homogenization, extrusion, or sonication, to disrupt any formed conglomerates.

## 4. Discussion

We have previously developed ultrasound-sensitive drug carriers that are safe for local drug release in NHPs [16, 17]. In the present work, we adapted the manufacturing process to enable efficient encapsulation and ultrasound-triggered release of hydrophobic antineoplastic agents. We demonstrated high encapsulation efficiencies for afatinib and doxorubicin and characterized their pressure-dependent release profiles in vitro. Together, these findings establish a platform for targeted chemotherapy delivery. The translational potential of this approach is further supported by a companion study employing the same microdroplet platform in a genetically engineered murine glioblastoma model [19] XXX.

This study shows that co-evaporation-based synthesis enables efficient encapsulation of highly hydrophobic antineoplastic agents into ultrasound-responsive microdroplets, addressing a major limitation of existing delivery systems. This method generates functional antineoplastic microdroplets that release the cargo upon the selective activation with focused ultrasound. This approach provides the necessary selectivity, which is crucial to avoid the systematic side effect associated with chemotherapy. Moreover, we have shown that the release is a function of the delivered energy. This provides the capacity to locally release these and other drugs at potentially much higher concentrations than previously possible, with release dose governed by the applied ultrasound pressure.

The drugs used in this study have been encapsulated using other methods. For instance, Pieper et al. reported doxorubicin encapsulation in PLGA particles at approximately 44% efficiency [10], and Missirlis et al. achieved up to 43% efficiency with PEG and poloxamer 407 [12]. However, neither of these methods provide ultrasound-gating. Similarly to these researchers, we found that the specific manufacturing technique used was key to achieving high encapsulation for these drugs.

The mechanism of ultrasound-triggered release is likely governed by a mechanical perturbation of the microdroplet structure [16, 17]. At sufficient acoustic pressures, the oscillatory motion and local pressure fluctuations may generate stress on the shell, promoting release of the encapsulated drug. In addition, acoustic diffusion and radiation forces can enhance mass transport and facilitate drug partitioning into the surrounding medium.

The polydispersity of the synthesized microdroplets presents an opportunity for further optimization. Although temperature is known to influence droplet size during formation, the mechanisms underlying the relatively larger and more heterogeneous size distribution observed with co-evaporation remain unclear. Future refinements of the fabrication process—such as the implementation of microfluidic synthesis or extrusion-based size control—may enable improved uniformity and tighter control over microdroplet size distributions.

In conclusion, we have developed an ultrasound-sensitive carrier that is capable of encapsulating antitumor agents, doxorubicin and afatinib, and releasing them upon the impact of focused ultrasound. We have previously validated the safety of the formulation in the brain of NHPs, and the approach is therefore poised for clinical translation. In sum, this study provides a platform for ultrasound-mediated, targeted chemotherapy.

## 5. Acknowledgements

The authors thank the members of The University of Utah Nanofab for assistance with scanning electron microscopy, and the members of the OneTarget Lab for helpful discussions and technical assistance. This work was supported by National Science Foundation (Grant No. CBET 2325125), the National Institutes of Health (Grant No. R61/R33), and the Huntsman Mental Health Institute and College of Engineering at the University of Utah. The content is solely the responsibility of the authors and does not necessarily represent the official views of the funding agencies.

## 6. Author contributions

Joshua Antonio Whiting: Conceptualization, investigation, methodology, formal analysis, visualization, writing – original draft.

Audri Yasmin Al Hasan Dara: Investigation, methodology, writing – review and editing.

James Kwan: Investigation, writing – review and editing.

Jan Kubanek: Conceptualization, supervision, resources, funding acquisition, writing – review and editing

