## Appendix A: Supplemental for "Development and Characterization of Ultrasound-Activated Polymeric Microdroplets for Targeted Chemotherapy"

### Appendix A. Supplementary

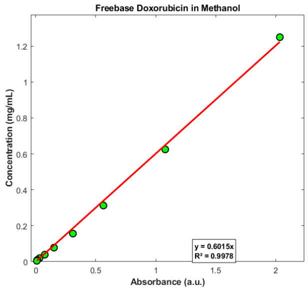


(a) DOX in methanol (encapsulation)

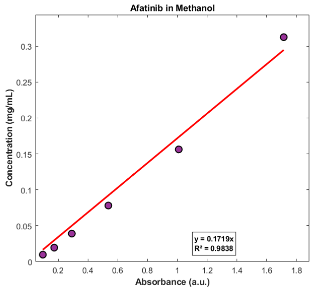


(b) Afatinib in methanol (encapsulation)

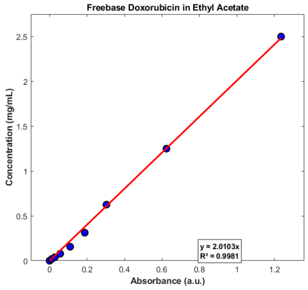


(c) DOX in ethyl acetate (release)

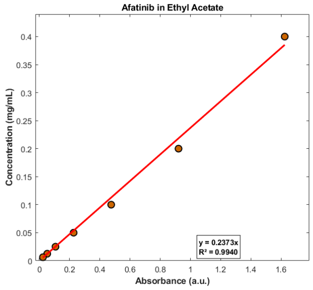


(d) Afatinib in ethyl acetate (release)

Figure A.5: UV–Vis standard curves used for quantification of encapsulation efficiency and ultrasound-triggered release. (a) Doxorubicin in methanol for encapsulation efficiency determination. (b) Afatinib in methanol for encapsulation efficiency. (c) Doxorubicin in ethyl acetate for release quantification. (d) Afatinib in ethyl acetate for release quantification. Linear regression on the methanol and ethyl acetate plots yielded $R^2$ values of 0.9978 and 0.9981 for DOX and 0.9838 and 0.9940 for afatinib, respectively.
